# Analysis of DNA transposition by DNA transposases in human cells

**DOI:** 10.1101/2023.04.26.538406

**Authors:** Yves Bigot, Makiko Yamada, Helen Mueller, Victor Morell, Sabine Alves, Thierry Lecomte, Alex Kentsis

**Affiliations:** PRC, UMR INRA0085, CNRS 7247, Centre INRAE Val de Loire, 37380, Nouzilly, France; Tow Center for Developmental Oncology and Sloan Kettering Institute, Memorial Sloan Kettering Cancer Center, New York, New York, USA; Department of Hepatogastroenterology and Digestive Oncology, Trousseau Hospital, Tours, France; Inserm UMR 1069, Nutrition, Croissance et Cancer, Université de Tours, Tours, France; Departments of Pediatrics, Pharmacology, and Physiology & Biophysics, Weill Medical College of Cornell University, New York, New York, USA

## Abstract

This manuscript discusses the recent report “Cognate restriction of transposition by *piggyBac*-like proteins” in *Nucleic Acids Research* by Beckermann et al and related recent publications about the inability to detect DNA transposition activity of human domesticated DNA transposase PGBD5. Measuring DNA transposition activity of transposases in human cells, where these enzymes can act on endogenous genomic substrates and induce DNA damage, is complicated by these and other cellular responses. Possible reasons for the discordant findings of Beckermann et al with prior independent reports of PGBD5 DNA transposition by Helou et al and Henssen et al and specific details of experimental methods in human cells are presented. In particular, by using independent experiments that reproduce PGBD5-mediated genomic integration, we demonstrate how supraphysiologic and ectopic overexpression of PGBD5 can cause DNA damage and cell death, and artifactual loss of apparent activity in clonogenic transposition reporter assays. While PGBD5 can support apparent DNA transposition, its cellular activity predominantly involves double-strand DNA breaks, deletions and other DNA rearrangements. We discuss the implications of this phenomenon for the interpretation of experimental assays and activities of domesticated DNA transposases.

## Introduction

*PGBD5* is the oldest known evolutionarily conserved domesticated DNA transposase gene in vertebrates (1,2). The presence of strong purifying selection and specifically restricted tissue expression for PGBD5 are analogous to RAG1/2, the second oldest evolutionarily conserved vertebrate DNA transposase. RAG1/2 is essential for somatic diversification of immunoglobulin receptor genes in developing immune lymphocytes in vertebrates (3). When dysregulated, RAG1/2 is also responsible for the induction of oncogenic chromosomal translocations in leukemias and lymphomas (4). The expression of PGBD5 in neuronal tissues and young-onset human solid tumors led to the hypothesis that site-specific nuclease activities of PGBD5 may contribute to normal neuronal development and human disease (5,6).

Previously, we found that human PGBD5, when expressed ectopically in human HEK293 cells but at near-physiologic levels similar to neuronal and human tumor cells, can mediate DNA transposition of synthetic episomal transposons into the human genome (2). This activity was defined using genomic PCR to measure and determine the location and sequence composition of the transposon integration breakpoints, as well as clonogenic fluorescence and antibiotic resistance reporter assays. PGBD5 activity required intact *piggyBac* inverted terminal repeat (ITR) transposons, including a G-rich trimer motif at its outer ends (8) that is important for the efficient enzymatic excision-insertion mechanism used by *piggyBac*-type DNA transposases. PGBD5 activity also required the presence of three specific aspartic acid residues in the transposase domain of PGBD5, two of which are conserved in other *piggyBac*-type DNA transposases. Most importantly, analysis of the human genomic integration breakpoints of transposons mobilized by human PGBD5 demonstrated a specific feature of the DNA transposition reaction by *piggyBac*-type DNA transposases (2), the use of a TTAA insertion site, at least at one of both ends. Subsequently, independent work using diverse DNA sources, cell lines and alternative methods in independent laboratories confirmed these findings (7,8). In particular, Helou et al found that PGBD5 can mediate both canonical DNA transposition in human cells, as well as alternative DNA integration reactions (7,8).

Importantly, studies of endogenous PGBD5 in human tumor cells revealed similar DNA transposition activity. Delivery of episomal *piggyBac* transposons via transfection of human G401 rhabdoid tumor cells that express PGBD5 endogenously was sufficient to mediate integrations into genomic DNA, including into genomic loci physically bound by endogenous PGBD5, as measured using chromatin immunoprecipitation and DNA sequencing (ChIP-seq) (9). This activity was completely abolished by the specific depletion of endogenous PGBD5 using RNA interference, as also independently observed in the Bigot laboratory (8). Lastly, studies of primary human RPE and BJ cells, in which PGBD5 is expressed ectopically at near-physiologic levels, revealed that PGBD5 is sufficient to induce human genomic rearrangements, leading to malignant cell transformation. This activity required the presence of the aspartate triad in PGBD5 (2), and cellular end-joining DNA repair, and involved specific human genomic sequences at the DNA rearrangement breakpoints, which were found to be enriched at somatic DNA rearrangements observed in human patient rhabdoid tumors that naturally express PGBD5. Importantly, ectopic expression of PGBD5 caused double-strand DNA breaks of human genomic substrates, which if unrepaired, led to the accumulation of DNA damage and cellular apoptosis (10). In all, these results demonstrated that human PGBD5 is active in human cells, mediating both canonical DNA transposition as well as alternative forms of DNA rearrangements due to its nuclease activity and other cellular effects, at least some of which can be pathogenic, causing cell death and cancerous transformation. These studies have raised many questions about the physiologic and pathophysiologic functions of domesticated DNA transposases and their molecular and cellular mechanisms (4).

We were therefore dismayed to read that Beckermann et al detected no significant PGBD5 transposition or integration activity in their recent paper published in *Nucleic Acids Research* (11). Here, we report additional complementary results that confirm DNA excision and genomic integration by human and mouse PGBD5. We provide several reasons for the discordant findings of Beckermann et al, including specific experimental details that are needed to detect the cellular activities of domesticated DNA transposases. While this manuscript was in preparation, Kolacsek et al also reported no detectable genomic integration activity of PGBD5 on transposon substrates in human cells (13). Here, we present an alternative analysis of their published experimental results, showing that PGBD5 has measurable cellular activity with a distinct substrate sequence preference as compared to the looper moth piggyBac. On balance, PGBD5 exhibits significant sequence-specific nuclease activity in human cells, as reproduced by multiple, diverse, and independent experimental systems and investigators. The extent to which PGBD5 retains ancestral piggyBac-like transposase activity remains to be defined using ongoing enzymatic biochemical and atomic resolution structural studies, which are required to establish the precise molecular mechanism(s) of this evolutionary conserved transposase-derived vertebrate gene.

## Materials and Methods

### Plasmids

Plasmid constructs were obtained as described (7,8). The plasmids pCS2-Mm523, pCS2-Mm409, pCS2-Hs554 and pCS2-Hs455 encode 5Xmyc-tagged PGBD5 isoforms that are 523, 409, 554 and 455 amino acid residues in length from mouse (Mm) or human (Hs) origins, respectively. Our ORF source of Hs455 was the plasmid pRecLV103-GFP-PGBD5 (Addgene plasmid 65409; RefSeq accession numbers NP_078830 and NM_024554). The ORF encoding Hs554 (corresponding to RefSeq and Uniprot accession numbers NP_001245240 and EAW69912, respectively) was synthesized by ATG:biosynthetics (Merzhausen, Germany). Mouse Mm523 and Mm409 orthologues correspond to Uniprot/RefSeq accession numbers D3YZI9.21/NP_741958.1 and D3YZI9.12/XP_006530867.1, respectively. The plasmids pBSK-IFP2-TIR5’-NeoR-TIR3’ (pBS-*Ifp2*-NeoR) and pBS-EF1-IRES-Neo-PB-ITR were used as transposon sources in excision and toxicity assays. pBSK-NeoR was a source of NeoR cassette void of any transposon sequences. pCS2-GFP was obtained from (12).

### Excision assays

Excision assays were performed as described (13,14). Briefly, for each sample, 100,000 HeLa cells in one well of 24-well plate were transfected with JetPEI (Polyplustransfection, Illkirch-Graffenstaden, France) using PEI nitrogen to DNA phosphate ratio of 5, and 220 or 400 ng pBS-*Ifp2*-NeoR and pCS2-transposase plasmid DNA at 1:0.1 or 1:1 ratio, respectively. Two days post-transfection, plasmid DNA was isolated using the Qiagen plasmid miniprep kit as described (14). DNA was quantified using the Qubit according to manufacturer’s instructions (Molecular Probes, Eugene, USA). The NeoR plasmid sequence was amplified using 5’-GGAGAGGCTATTCGGCTATG-3’ and 5’-GGTGAGATGACAGGAGATCC-3’ primers. The transposase DNA sequences were amplified using 5’-AGCATGTTCTGCTTCGACG-3’ and 5’-AGGAAACAGCTATGACCATG-3’ primers in the pCS2 plasmid. For plasmids encoding each of the four PGBD5 isoforms, PCR was performed using 5’-GAGCTGGAAGAAGTCCACCG-3’ primer for Mm523 and Mm409; 5’-GGCTTATTCTTCAGCACATCC-3’ for Hs455; 5’-CTGAAATACTTCCACGTGGTG-3’ for Hs554, and 5’-TAATACGACTCACTATAGGGCG-3’ primer in the pCS2 plasmid. PCR was performed in a 50 µL reaction volume with ten nanograms of template DNA, 0.3 U GoTaqG2 Flexi (Promega, Madison, WI), 2.5 mM MgCl_2_, 200 µM dNTP and 100 nM of each primer. PCR was carried out for 30 cycles (94°C for 30 sec, 52°C for 30 sec, 72°C for 30 sec). To detect excision products, ten nanograms of purified DNA was used in a two-round nested PCR assay in which primers were specific to the plasmid backbone and resulted in a PCR product only from the excised and repaired plasmid copies due to large transposon cassettes. First-round PCR primers were as follows: 5’-CCCAGTCACGACGTTGTAAAACG-3’, 5’-AGCGGATAACAATTTCACACAGG-3’. Thirty-five cycles were done in a 50 µL reaction volume with 0.3 U GoTaqG2 Flexi (Promega, Madison, WI), 2.5 mM MgCl_2_, 200 µM dNTP and 100 nM of each primer. The first-round PCR program was for 35 cycles: 94°C for 30 sec, 62°C for 30 sec, 72°C for 30 sec. Second-round PCR primers were as follows: 5’-CGTTGTAAAACGACGGCCA-3’, 5’-AGGAAACAGCTATGACCATG-3’. Thirty-five cycles were performed under similar amplification conditions using 1 µL each of first-round PCR products, and an annealing step at 55°C. PCR products were purified using Nucleospin kit according to manufacturer’s instructions (Macherey-Nagel, Düren, Germany), and visualized by electrophoresis using 2.5 % LE agarose gel. To sequence excision and non-specific products, PCR products were excised from gels, purified using Nucleospin kit (Macherey-Nagel, Düren, Germany), and subcloned into pGEM-T plasmid, followed by Sanger sequencing of both strands using second-round PCR primers. Assays were performed in triplicate and experiments were replicated three times.

### Integration assays in HeLa cells

Assays were performed as described (12). Briefly, for each sample, 100,000 HeLa cells in one well of 24-well plate were transfected with JetPEI (Polyplus-transfection, Illkirch-Graffenstaden) and 400 ng DNA plasmid and with or without equal amount of donor NeoR plasmid. Two days post-transfection, cells were trypsinized, replated in 100 mm dish, and incubated in culture medium containing 800 μg/mL G418 sulfate (Eurobio Scientific, Les Ulis) for 15 days. Plates were washed twice with phosphate buffered saline (PBS), fixed and stained overnight with 70% EtOH and 0.5% methylene blue, and colonies > 0.5 mm in diameter were counted. Experiments were performed in triplicate and replicate twice.

### Integration assays using HEK293 cells

HEK293 cells were obtained from ATCC (Manassas, VA, USA), and used at passage 4-7 after plating. Sixty thousand HEK293 cells per one well of 24-well plate were plated one day before transfection. All the assays (n=7) were performed in triplicate. Thirty minutes before transfection, the media was changed into antibiotics-free DMEM with 10% fetal bovine serum (FBS), 1% L-glutamine, and 1% sodium pyruvate. The transfection complex was made in 50 µl of Opti-MEM by adding 250 ng of transposon and transposase plasmids respectively, followed by addition of 1 µl of FuGene6 (Promega, Madison, WI) and incubated for 15 min. PB-EF1-MCS-IRES-NEO (System Biosciences, Mountain View, CA) was used as a transposon and its inverted terminal repeats-deleted version was used as a negative control transposon in combination with either pRecLV103-GFP, pRecLV103-GFP-PGBD5, or T. ni piggyBac transposase (PB200PA-1; System Biosciences) (2). Sequences of all the five plasmids were validated by whole-plasmid sequencing using Primordium (Monrovia, CA). Fourteen to eighteen hours later, the media was completely changed into DMEM with 10% fetal bovine serum, 1% L-glutamine, 1% sodium pyruvate, and 1% penicillin/streptomycin. After 48 hours of transfection, cells were trypsinized and cell viability was quantified using trypan blue dye exclusion, with 30,000 viable cells replated on 10-cm dishes in media supplemented with 1 mg/ml G418 sulfate (Gibco, Billings, MT). After two weeks of selection with G418, plates were washed in PBS and fixed with 4% paraformaldehyde/ 0.1M phosphate buffer for 15 min. After washing in PBS, cells were stained with Crystal violet and colonies were counted. Colony was defined as a cell cluster consisting of more than 20 cells and the colony number was counted using a benchtop brightfield microscope. For the re-analysis of the transposition assays conducted by Kolacsek et al (15), a high-resolution image of Figure 2B was downloaded and the colony numbers for MER85 and PB-puro were independently counted by three observers. The mean of duplicates from three observers were plotted.

### Protein expression analysis

For each sample, 100,000 HeLa cells in one well of 24-wells plate were transfected with JetPEI (Polyplustransfection, Illkirch-Graffenstaden, France) using PEI to DNA ratio of 5, and 400 ng of plasmid DNA encoding GFP or PGBD5 variants. Forty-eight hours post transfection, adherent cells were washed twice with phosphate buffered saline. Cells were lysed and proteins were extracted by vigorous pipetting with 300 µL of Laemlli buffer containing 0.1% 2-mercaptoethanol at room temperature (RT), sonicated twice for 5 min at 50W, 46 kHz using the Ultrasonic Jewelry Cleaner Leo (Leo, New Taipei City, Taiwan). Lysates were then heated at 95°C for 10 min, cooled for 5 min at RT and centrifuged for 1 min at 15,000 *g* at RT. Lysates cleared by centrifugation (20 µL) were resolved by electrophoresis using 4-20% mini-Protean TGX gels in Tris/Glycine/SDS buffer (Bio-Rad, Hercules, USA). Gels were then transferred onto 0.2 µm nitrocellulose membranes using Trans-Blot Turbo Transfer System gel (Bio-Rad, Hercules, CA, USA). Membranes were incubated overnight at 4°C in Odyssey blocking buffer containing 0.1% Tween-20 at 4°C and 1/1000 dilution of anti-c-myc mouse monoclonal antibody (Catalogue number 11 667 149 007, Roche, Bâle, Switzerland) or anti-α-tubulin mouse antibody (Catalogue number T9026, Sigma, Saint-Louis, MI, USA). Membranes were washed three times with phosphate buffered saline containing 0.1% Tween-20 at RT, and incubated for 1 hour at RT in Odyssey blocking buffer containing 0.1% Tween-20 and 1/1000 dilutions of anti-mouse secondary antibody conjugated to IR-800 and anti-rabbit secondary antibody conjugated to IR-670. Membranes were washed three times with phosphate buffered saline containing 0.1% Tween-20 at RT, and imaged using Odyssey scanner (LICOR Biosciences, Lincoln, NE, USA). Experiments were repeated five times.

### Toxicity assay in HEK293 cells

One hundred thousand HEK293 cells (ATCC, Manassas, VA, USA) were plated in one well of 24-well plate and transfected using 0.5 µl PEIPro (Polyplus, Strasbourg, France) with 500 ng of total plasmid DNA, containing 250 ng of pBS-EF1-IRES-Neo-PB-ITR and 250 ng of either pRecLV103-GFP or pRecLV103-GFP-PGBD5, respectively. Two days after transfection, green fluorescent cells were sorted using BD FACSAria cell sorter (Franklin Lakes, NJ, USA) and replated. GFP-positive cells were divided into low and high populations at a ratio of 1:1. Batches of 12,000 cells of each population were seeded in one well of 12-well plates and grown in DMEM with 10% fetal bovine serum, 1% L-glutamine, and 1% penicillin/streptomycin for 10 days.

### Statistical analyses

Analyses were performed using Prism 6 (GraphPad Software Inc, San Diego, CA, USA) and OriginPro (OriginLab, Northampton, MA, USA). All t-tests were two-tailed. F tests were used to compare variances of samples and Welch’s correction was used when they were significantly different (*p*<0.05).

## Results

Previously, both Bigot and Kentsis groups and Beckermann et al (11) used clonogenic assays with plasmids encoding neomycin resistance genes flanked by *piggyBac* (*Ifp2*) transposon ITRs to assay their excision and genomic transposition upon co-expression of transposases. These assays use transient transfection of transposase-expressing cells followed by antibiotic selection to estimate genomic integration of antibiotic resistance-conferring transposons. Combined with the DNA sequencing analysis of episomal transposon excision and genomic integration sites, these assays can be used to accurately define and quantify genomic DNA transposition. While Beckermann, Luo, Wilson and colleagues performed extensive experiments using varied experimental conditions, including replications of our published experiments, several of their experimental conditions were nonetheless apparently different. For example, variability in the levels of ectopic DNA transposase expression is an important determinant of cellular activity. With too low expression levels, no excision and no integration by transposition and recombination can be observed. On the other hand, high expression levels can lead to unrepaired DNA damage, cell mitotic arrest, cell death or even a low or null transposase activity due to its overexpression, which confound the interpretation of clonogenic assays, and at sufficiently high levels, abrogate measurements of genomic integration entirely.

To examine these issues directly, we used the cDNA sequence encoding human PGBD5 as originally reported (2), which was also called PGBD5v1 in (11) and which we also term Hs455, as well as its alternative longer isoform called Hs554. We also included the corresponding 409 and 523 amino acid mouse isoforms, which we called Mm409 and Mm523; the human isoform orthologue, Hs524, is called PGBD5v2 (11). Each of these isoforms was cloned into pCS2 DNA plasmids encoding Myc-tagged transposases; pCS2 is similar in size and sequence to the pCMV plasmid used in (11). Transient transfection of HeLa cells with equal amounts of plasmid DNA encoding Myc-tagged transposases revealed varying levels of protein expression as assayed by Western blotting and measured 2 days after transfection, Mm523>Hs455>Mm409>Hs554, consistent with variable cellular stability of different PGBD5 isoforms (Figure 1A).

**Figure 1.**
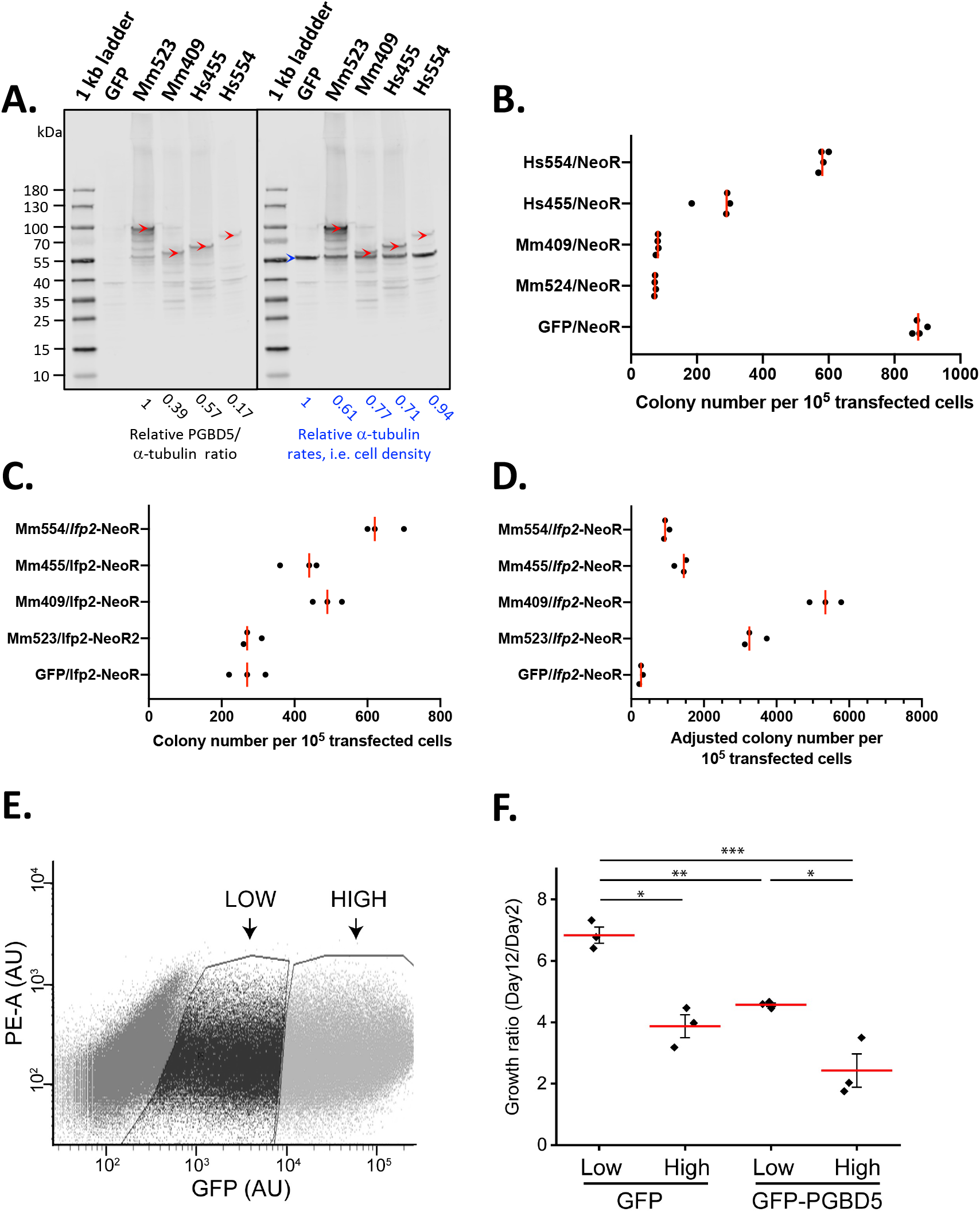
Ectopic PGBD5 overexpression induces cytotoxicity and impairs clonogenic capacity of human cells. (**A**) Western blot analysis of PGBD5 isoform expression in HeLa transfected with pCS2-GFP (lanes 2 and 8), -5Xmyc-Mm523 (lanes 3 and 9), -Mm409 (lanes 4 and 10), -Hs455 (lanes 5 and 11), and -Hs554 (lanes 6 and 12). The molecular weight markers are shown in lanes 1 and 7. Lane 1 to 6, the blot was probed with anti-myc mouse monoclonal antibody. Lane 7 to 12, same blot was probed again using anti-α-tubulin mouse monoclonal antibody. The relative PGBD5/α-tubulin ratios were calculated using image densitometry from five replicates. Red arrowheads mark the intact 5Xmyc-PGBD5 isoforms and the blue arrowhead marks α-tubulin. The apparent molecular weight of intact 5Xmyc-PGBD5 isoforms was higher than expected from their sequences: Mm523, 95-100 versus 64 kDa; Mm409, ∼60 versus 53 kDa; Hs455, 65-70 versus 58 kDa; Hs554, 80-85 versus 64 kDa, respectively. The presence of discrete bands of lower molecular weights also indicated that each of the four transfected PGBD5 isoforms was partly proteolyzed in cells. (**B**) Rates of integration of a NeoR cassette when recombination was mediated by GFP, Mm523, Mm409, Hs455 and Hs554, as normalized to GFP controls. (**C**) Rates of integration of *Ifp2*-NeoR when recombination was mediated by GFP, Mm523, Mm409, Hs455 and Hs554. (**D**) Rates of NeoR clones resulting from the integration of *Ifp2*-NeoR when recombination was mediated by GFP, Mm523, Mm409, Hs455 and corrected by the impairment of NeoR clonogenic capacity as measured in (B). Median values are marked by red lines. (**E**) Representative FACS plot of GFP-low and GFP-high cells, as indicated by arrows. (**F**) Growth of cells expressing high and low GFP or GFP-PGBD5, as assessed on day 12 relative to day 2 of culture; *p<0.05, **p<0.01, ***p<0.005.

Variability in a transposase expression is an important determinant of its transposition activity. Similarly, clonogenic assays with antibiotic reporters depend on the proliferation, survival and antibiotic tolerance of cells over multiple cell divisions. To assess these effects, we used a plasmid that encodes neomycin resistance but lacks transposon ITRs which are required for excision and transposition (Figure 1B). As expected, we observed that in the absence of any episomal transposons, human and mouse PGBD5 transposase expression was associated with significant impairments of clonogenic capacity and ultimate neomycin resistance, as compared to control HeLa cells transfected with GFP control plasmids (approximately 12-, 11-, 3-, and 2-fold for Mm523, Mm409, Hs455, and Hs554, respectively; Figure 1B). This PGBD5-induced impairment of clonogenic capacity was strongly correlated with the PGBD5 isoform expression levels, as quantified using image densitometry of Western immunoblots (*r*^*2*^ = 0.95; Figure 1A). This cellular toxicity is directly linked to the PGBD5-mediated induction of double-strand DNA breaks in the human genome, which requires active end-joining DNA repair, as cells deficient in specific forms of end-joining DNA repair undergo apoptosis or irreversible mitotic arrest, specifically when expressing wild-type PGBD5, but not its catalytically deficient aspartate-to-alanine mutant (10).

We then repeated this experiment using plasmids containing a neomycin resistance gene flanked by 345- and 252-bp from the 5’ and 3’ terminal sequences of the *piggyBac* transposon used previously (7,8) and shown in Figure 1C-D. When adjusted for the PGBD5-induced reduction in clonogenic capacity, we found that cells transfected with Mm523, Mm409, Hs455, and Hs554 PGBD5 isoforms exhibited approximately 12-, 20-, 4- and 4-fold greater numbers of neomycin-resistant colonies, as compared to control GFP-transfected cells (Figure 1C-D). We note that HeLa cells are known to have a relatively high spontaneous DNA integration rate due to their intrinsic genomic instability (16). In an independent experiment, we transfected plasmids encoding GFP-PGBD5 fusion protein and used fluorescence-activated cell sorting (FACS) to isolate cells expressing high versus low expression of GFP-PGBD5 (Figure 1E). Cells expressing high levels of GFP-PGBD5 exhibited significantly impaired proliferation and survival, as compared to low expressing and control cells (Figure 1F). These results recapitulate our previously published observations, both with respect to the PGBD5-induced genomic transposon integration and its cellular toxicity (2,7,8,10). Thus, detection of PGBD5-mediated genomic integration activity on synthetic transposon substrates requires optimization of the level of transposase expression when expressed ectopically in heterologous cells. For cytotoxic and mutagenic factors such as human PGBD5, the range of experimental conditions for observing its genomic activity in human cells is indeed quite narrow, which is different from the activity of heterologous enzymes such as piggyBac from the *T. ni* looper moth.

In order to confirm that the observed genomic integration of neomycin resistance transposon reporters was due to DNA transposition or other mechanisms of genomic DNA integration, we isolated plasmid DNA containing *piggyBac* transposons from transfected cells to measure ITR sequence-specific transposon excision. In Beckerman et al (11), these excision assays were performed using plasmid DNA samples purified from cells 24 hours post transfection. The experimental protocol was described as being identical to those used in (2) and two prior papers from the Wilson laboratory (17,18). However, in prior published studies, plasmid DNA was isolated 48 and 72 hours post-transfection in order to accumulate sufficient amounts of excision products in transfected cells. Similar longer duration was used in (14). Liu et al (13) and Wilson et al (19), utilized even longer durations of 72 and 96 hours. Similar observations have been made for other DNA transposases as well. For example, post-transfection duration of 18-30 hours was inadequate for the detection of excision products by the Sleeping Beauty DNA transposase, instead requiring 45-89 hours for adequate detection (13).

To examine this issue directly with PGBD5, we performed new excision assays using plasmid DNA purified from cells transfected with plasmids encoding PGBD5 and the same *piggyBac* transposon donor, and analyzed cells at 48 hours after transfection using PCR. We observed the specific 148-bp excision product in cells transfected with PGBD5 isoforms but not GFP control, corresponding to the canonically transposase-excised fragment with TTAA breakpoints (red arrowhead, Figure 2A-B). Insect *piggyBac* served as additional control, exhibiting an identical excision activity (Figure 2A). We also detected additional products, corresponding to non-canonical excision products with an alternative TTAA breakpoint (blue arrowhead, Figure 2A and 2C), as well non-specific amplicons (black arrowhead, Figure 2A). Detection of transposase excision products by PCR requires optimization of annealing and extension parameters, which was extensively done by Henssen et al to preferentially detect canonically excised products (2).

**Figure 2.**
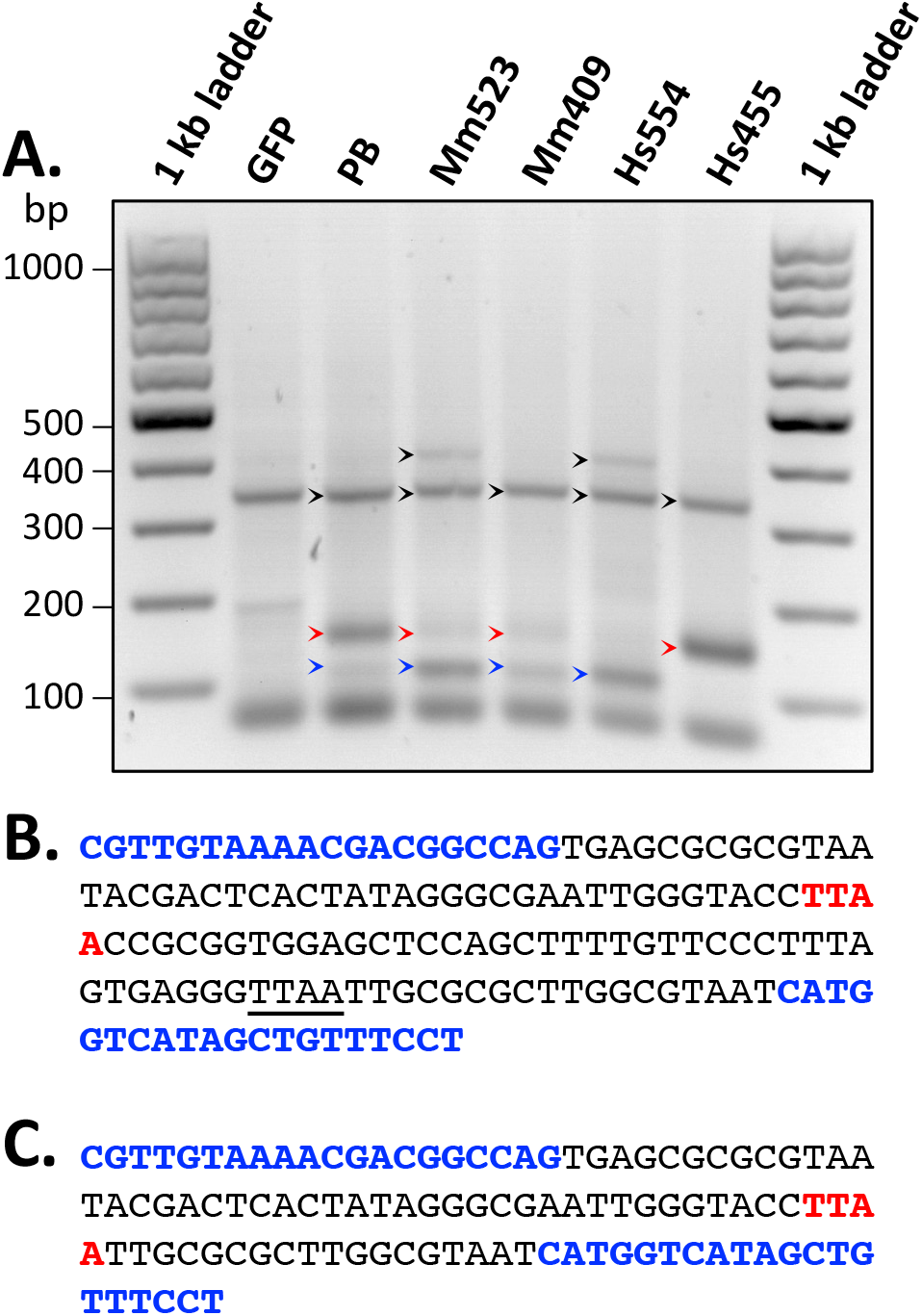
Human and mouse PGBD5 isoforms exhibit both canonical and non-canonical excision of episomal *piggyBac* inverted terminal repeat transposons. (**A**) Excision assays using pCS2-GFP (lane 2), -Mm523 (lane 3), -Mm409 (lane 4), -Hs455 (lane 5), -Hs554 (lane 6) or -PB (lane 7) in HeLa cells transfected with pBS-*Ifp2*-NeoR. The expected 148-bp excision products are marked by red arrowheads. Non-canonical excision products with an alternative TTAA breakpoint are marked by blue arrowheads, and non-specific amplicons by black arrowheads. (**B**) DNA sequence of the 148-bp excision product, as obtained by Sanger sequencing. (**C**) DNA sequence of the 105-bp non-canonical excision products. The non-canonical TTAA is underlined in (**B**). In (**B**) and (**C**), the restored TTAA motif after excision is marked in red, and second-round PCR primers are marked in blue.

However, this does not mean that PGBD5 functions exclusively or physiologically as a DNA transposase. Published work demonstrated that in addition to canonical DNA transposition of synthetic *piggyBac* transposons, PGBD5 also induces double-strand breaks on human genomic substrates, which are associated with deletions, inversions, and translocations (9). Our current results demonstrate that these non-transposition DNA reactions can also occur on episomal substrates (Figure 2A). However, detection of PGBD5 DNA transposition activity as assessed by the excision of episomal substrates requires sufficiently long (at least 48 hours post transfection) expression in cells, as also documented for other DNA transposases (vide supra).

In addition to replicating PGBD5-mediated genomic transposon integration in HeLa cells by Helou et al (7,8), as shown in Figure 1, we also sought to replicate the original observations of Henssen et al in HEK293 cells (2,9). We used newly obtained stocks of HEK293 cells and transfected them using plasmids encoding GFP-PGBD5 and neomycin resistance gene flanked by *piggyBac* ITRs, with *T. ni* piggyBac as positive control (Figure 3A). To assess *piggyBac* ITR-mediated transposition versus other forms of genomic transposon integration, we used identical plasmids encoding neomycin resistance gene with deleted ITRs (Figure 3A). We controlled for variable cytotoxicity by plating equal numbers of viable cells, as assessed after 2 days of transfection, and estimated transpositional capacity by calculating the ratio of the number of neomycin resistant colonies of cells transfected with intact versus deleted ITR plasmids (Figure 3B). While we observed significantly increased transposon genomic integration in cells expressing GFP-PGBD5 versus GFP (mean 1.3-fold; t-test *p* = 0.025; Figure 3B), this effect was substantially less than our original measurements (2,9). This may be indeed due to the overestimation of PGBD5 genomic integration in our prior studies, differences in expression rate due to transfection efficiency, or alternatively, to differences in host cell factors which are known vary in different clones of HEK293 cells (20).

**Figure 3.**
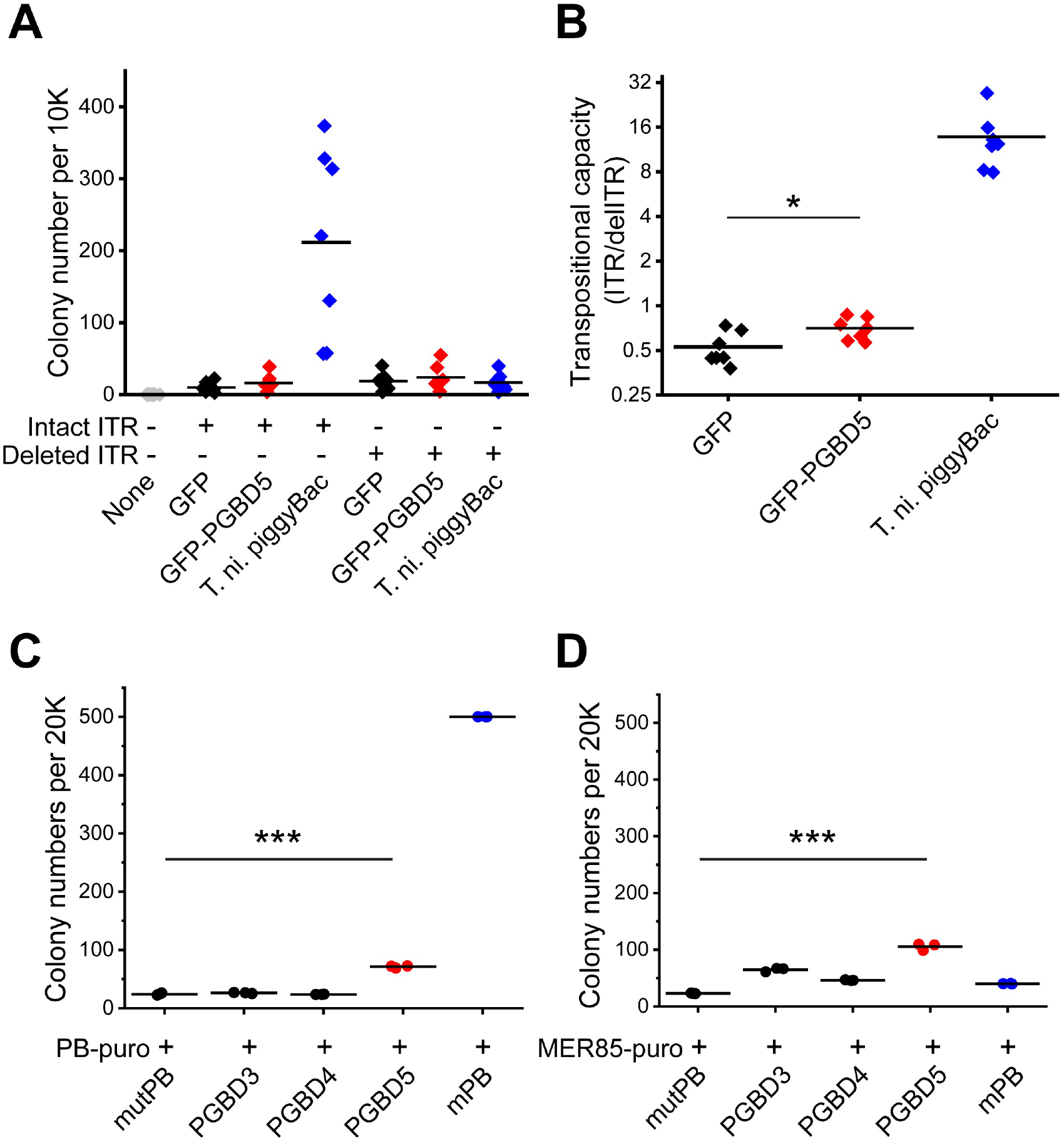
Ectopic PGBD5 overexpression in HEK293 cells induces transposition of *piggyBac* ITR transposons with relatively low efficiency as compared to *T. ni* piggyBac. (**A**) Quantification of raw colony numbers of Neomycin-resistant clonogenic assay using GFP (negative control in black), GFP-PGBD5 (in red), and *T. ni*. piggyBac (positive control in blue), in combination of either the NeoR reporter flanked by the *piggyBac* inverted terminal repeats (PB-EF1-MCS-IRES-NEO carrying the intact ITRs: abbreviated as Intact ITR) or control reporter lacking ITRs (PB-EF1-MCS-IRES-NEO without ITRs: abbreviated as Del-ITR). Bars indicate mean values of 7 biological replicates, performed by 3 blinded independent experimental operators. (**B**) Transpositional capacity, i.e., ratio of colony numbers obtained using the Intact ITR to the colony numbers obtained by the Del-ITR (GFP vs GFP-PGBD5; *n* = 7; two-tailed t-test *p* = 0.025*). (**C** and **D**) Re-analysis of the transposition assays conducted by Kolacsek et al 2022 (15). Either a puromycin cassette flanked by the *piggyBac* terminal inverted repeats (**C**) or by the *MER85* terminal inverted repeats (**D**) were co-transfected with mutant piggyBac transposase (mutPB), PGBD3, PGBD4, PGBD5, and piggyBac transposase (mPB). Three independent observers counted colony numbers using the images of Fig. 2B from Kolacsek et al 2022. Each symbol represents the mean count of duplicates from a single observer. Bars indicate mean values of three observers. PGBD5 in red shows more colony numbers than mutPB (two-tailed t-test \*\*\**p* = 6.0e-6 for (**C**) and \*\*\**p* = 2.0e-5 for (**D**)).

While this work was in progress, Kolacsek et al also reported measurements of PGBD5 genomic integration activity of transposon substrates in HEK293 cells (15). Fortunately, their publication included high-resolution photographs of antibiotic resistance clonogenic reporter assays for PGBD5 and other transposase-derived genes PGBD3 and PGBD4, as compared to the insect piggyBac transposase. We used three blinded observers to re-quantify the published photographs. Contrary to the report of Kolacsek et al, Orbán and colleagues (15), we observed significant activity of PGBD5 but not PGBD3 and PGBD4 on insect *piggyBac* transposons (t-test *p* = 6.0e-6 of PGBD5 versus catalytically inactivate piggyBac, mutPB; Figure 3C). Intriguingly, PGBD5 also exhibited significant genomic integration activity of synthetic transposons containing human MER85 ITRs (Figure 3D). In all, these results indicate that PGBD5 retains measurable sequence-specific genomic integration activity on piggyBac-like transposons.

Finally, Beckermann et al (11) used two additional reporter assays to measure the cellular activity of PGBD5 to mobilize synthetic transposons into episomal plasmids (antibiotic plasmid rescue assay) and to be recruited to synthetic ITRs using a transcriptional activator reporter (luciferase assay). These are widely used assays to measure the activity of autonomous transposases, such as the insect *piggyBac*. However, PGBD5 exhibits relatively low cellular activity as compared to insect piggyBac, likely due to the evolutionary adaptations during its vertebrate domestication. Thus, cytotoxicity of ectopic PGBD5 overexpression, combined with its relatively low cellular activity, requires highly sensitive assays. This is further compounded by the apparent requirement for cellular PGBD5 cofactors, which underlie differences in behavior of endogenous PGBD5 (such as in human tumor cells and neurons) as compared to ectopically expressed PGBD5 in heterologous cells, which lack the relevant PGBD5 cofactors and DNA repair functions. In contrast to the insect piggyBac which is an autonomous mobile (parasitic) genetic element, PGBD5 is a domesticated transposase, evolved for physiologic functions on chromatin substrates in vertebrate cells.

Thus, Helou et al (7,8), and Henssen et al (2,9) measured genomic DNA transposition of human and mouse PGBD5 isoforms into the human genome using LAM-PCR and FLEA-PCR, respectively. Beckermann et al (11) noted that both assays rely on PCR amplification with its associated potential for artifacts and false positive results. Indeed, all assays have imperfect sensitivity and specificity, requiring proper controls to establish them directly. For example, one of the many controls used by Henssen et al (2,9) includes mutant ITR transposons in which the essential 5′-<u>GGG</u>TTAA-3′ hairpin structure was mutated to 5′-<u>ATA</u>TTAA-3′ (location of mutation underlined). Transfection of the intact ITR led to the identification of unique transposon integration sites with TTAA breakpoints, consistent with apparent DNA transposition, in contrast to the mutant ITR, which produced significantly reduced and largely non-canonical integration sequences lacking TTAA junctions (2). In fact, this assay has sufficient sensitivity and specificity to detect DNA transposition upon transfection of the GGG-intact as compared to the ATA-mutant transposon donors even in human G401 rhabdoid tumor cells which express relatively low levels of endogenous PGBD5 (9). Cells depleted of endogenous PGBD5 using specific RNA interference are consistently unable to carry out genomic transposon integration (8). Indeed, the requirement of sensitive cellular assays to observe PGBD5 activity is consistent with the relatively low genomic integration activity of PGBD5 (2). In fact, the *piggyBac* ITR transposon-specific activity of PGBD5, as compared to other forms of genomic integration, is near the limit of detection using current clonogenic assays, as can be seen from our new replication experiments in HeLa and HEK293 cells (Figures 1 and 3A-B), as well as re-analysis of published results (15; Figure 3C-D).

Interestingly, the analysis of Kolacsek et al took into account the impact of transposase expression on cell viability. To do this, cell death was assessed using propidium iodide uptake and flow cytometry (21), and 48 hours post transfection cells were washed to remove cell culture medium. However, this procedure would also remove dead detached cells, as collected by trypsinization and centrifugation. Thus, the measurements of viable propidium iodide-negative cells are not appropriate (22,23). For this reason, we prefer to assess cytotoxicity effects over the whole time of the assay, involving transfection, replating and antibiotic selection (Figure 1B). Indeed, transient expression of transposases such as piggyBac can extend for as long as 7 to 10 days post-transfection (12,24).

Lastly, Beckermann et al (9) suggested that the PGBD5-induced DNA transposition as assessed by LAM-PCR by Helou et al (7,8) and using FLEA-PCR by Henssen et al were not consistent with each other. To address this, we independently reproduced the analysis of these studies. All data are publicly available via NCBI Sequence Read Archive as listed in their original publications, as well as directly accessible from https://doi.org/10.5281/zenodo.3737880. We identified genomic integration events by mapping reads with unique split sequences between the human genome and the *piggyBac* transposons. Henssen et al used a stringent threshold of at least 20 unique split reads to quantify genomic transposition by human PGBD5. This identified 65 canonical and 14 non-canonical integration sites. Reducing this threshold to 4 unique reads identifies 131 and 145 canonical and non-canonical sites, respectively (Figure 4A-D). Reducing this threshold to 2 unique reads to maximize sensitivity, which was the relatively permissive parameter originally used by Helou et al, produces 131 canonical and 297 non-canonical sites, which are approximately equal to the rates of canonical and non-canonical genomic transposition originally reported by Helou et al for mouse Pgbd5 (Figure 4A-D). Thus, mammalian PGBD5 can mediate DNA integration events of synthetic *piggyBac* transposons with a canonical insertion signature by transposition present at least at one of both ends, as well as non-canonical DNA rearrangements, as reported by multiple studies carried out independently in different laboratories with diverse reagents and assays, and now reproduced again here. In contrast to the insect *piggyBac* which supports mostly precise genomic insertions of its transposons, most of the PGBD5-mediated insertions are imprecise (Figure 4E). As recently highlighted for transposition assays of the piggyBac and Sleeping Beauty transposases, their transfection maintains expression for 7-10 days (24), PGBD5 is also likely present for a similar duration in cells (Figure 1E-F). So, the resulting integration and genomic remodeling profile is likely the product of several rounds of potentially tiled events triggered by PGBD5, its different DNA binding sites, and the DNA repair machinery (9).

**Figure 4.**
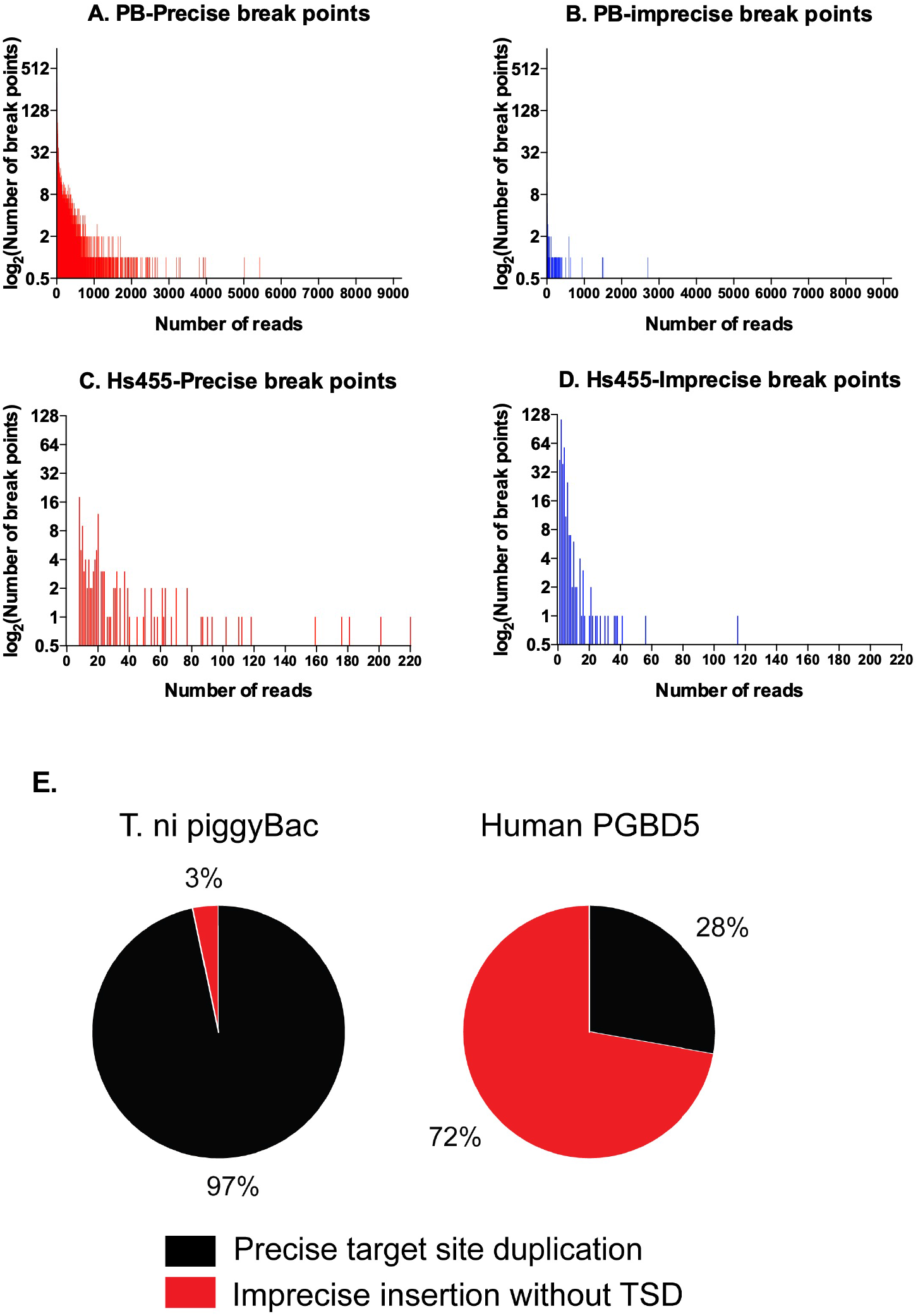
PGBD5 induces both canonical human genomic transposition and non-canonical DNA integration in human cells. Graphic representations of the numbers of obtained reads among *Ifp2* break points when PB (**A** and **B**) and Hs455 (**C** and **D**) mediated integration events. (**A**) and **(C**) Distribution of precise break points. (**B**) and **(D**) Distribution of imprecise break points. In (**A**) and **(B**), the data were extracted from (7) and (**C**) and **(D**) from (2). (**E**) Pie charts showing the ratios of the total number of precise break points (in black, obtained from (**A**) and (**C**)) and total numbers of imprecise break points (in red, obtained from (**B**) and (**D**)) for *T. ni* piggyBac (left) and PGBD5 (right), respectively.

## Discussion

Is PGBD5 a DNA transposase? Multiple studies have shown that *PGBD5* is a vertebrate transposase-derived gene, distinct from the *piggyBac* transposases in other organisms (1,8,9). As evident from its chromosomal immobilization and purifying selection in vertebrates, human *PGBD5* has evolved away from being an autonomous mobile genetic element (in contrast to *Trichoplusia ni piggyBac* and *Myotis lucifugus piggyBat*) and has become domesticated as an essential gene with endogenous physiologic functions. As such, *PGBD5* has co-evolved with its cellular cofactors and its divergent RNase H-like transposase domain to carry out evolutionarily selected functions in neurons, where it is expressed nearly exclusively in mammals. The physiologic functions of PGBD5 are under active investigation in multiple laboratories, including our own. Our recent work indicates that PGBD5 is required for normal human and mouse brain development due at least in part to its nuclease activity and associated with recurrent somatic DNA deletions (Jubierre et al. 2023, *bioRxiv*). These and future studies should reveal how the enzymatic activities of the PGBD5 transposase domain contribute to normal mammalian neuronal and brain development, and define its endogenous physiologic substrates. Likewise, PGBD5 is also expressed in the majority of solid tumors in children and young adults. Studies to date have shown that its dysregulation promotes the induction of sequence-specific oncogenic DNA rearrangements, nominating PGBD5 as the long sought-after developmental mutator for these cancers (4-6). Given these considerations, it is unlikely that mammalian PGBD5 functions physiologically as a DNA transposase per se.

Indeed, while PGBD5 retains residual ability to carry out apparent transposition of synthetic episomal *piggyBac* transposons in human cells, observations of PGBD5 activity on human genomic substrates have revealed deletions, inversions, translocations and other DNA rearrangements, without any evidence of “cut-and-paste” transposition thus far. This is analogous to the domesticated vertebrate DNA transposase RAG1/2, which can induce translocations and other DNA rearrangements when dysregulated in cancer cells (25). RAG1/2 mediates physiologic somatic deletions in lymphocytes, but can also be engineered to carry out DNA transposition of synthetic substrates (26,27). Ongoing biochemical and structural studies should reveal physiologic substrates, cellular cofactors, and mechanisms of DNA recognition and enzymatic remodeling by PGBD5.

We agree with Beckermann et al (11) that lepidopteran piggyBac transposase has specific activity on its *piggyBac* transposons. In contrast to human PGBD5, insect piggyBac failed to detectably mobilize synthetic transposons containing human PGBD5-specific signal sequences (PSS). However, it was recently described that the lepidopteran piggyBac transposase is able to mobilize related piggyBac-like elements (PLE) (28), but not the distantly related piggyBat element, which in addition displays a fully different sequence organization at its ends (11). The ability of lepidopteran piggyBac to bind to numerous non-PLE chromosomal DNA sites (29) also suggests that it might bind to chromosomal DNA motifs displaying features of PLE-like ends (30). We also agree that in addition to apparent DNA transposition human PGBD5 can mediate double-strand breaks and other DNA rearrangements, presumably because it probably lacks a sequence-specific high-affinity DNA binding domain, which is conferred by the CRD domain in the insect piggyBac transposase.

However, “cognate restriction of transposition” by DNA transposases is not a discrete phenomenon. Rather, cellular activities of transposases and nucleases are relative activities dependent on enzyme concentration, cellular factors, and DNA substrates. This is well established for restriction endonucleases for example, where they bind and cleave both higher-affinity (specific) and lower-affinity sequences to varying degree, as determined by relative binding affinities, catalytic activities, and solution conditions. Ectopic expression of PGBD5 can transform human cells due to the induction of oncogenic DNA rearrangements. Recent studies of the clinical use of engineered piggyBac DNA transposase for therapeutic gene transfer have also raised questions about its cellular substrates, and the possibility of inducing human genomic rearrangements (29). We hope that future collaborative research will continue to define the specific mechanisms and functions of these semantically simple, but biologically complex molecules.

## Acknowledgements

We thank members of our groups for critical feedback.

